# Fracture Analysis of Repaired Damage Induced Calcium Sulfate Dihydrate

**DOI:** 10.64898/2026.05.27.728004

**Authors:** Nestor Alvarez-Rodriguez, Hobbs McAllister, Alexander Lee, Brock Guillot, Luke Soileau, Kevin Hoffseth

## Abstract

Bone fragility is a major ang growing public health concern for many with degenerative bone diseases. A better understanding of fracture propagation in bone can better inform treatment options for these individuals. Human bone is an anisotropic material, with directional microstructure that fractures can propagate around. In a healthy individual, the bone remodeling process repairs damaged tissue. Calcium sulfate dihydrate is a well established surrogate material for bone. This study will mechanically investigate damage repaired plaster samples using three point bending to characterize crack propagation around repaired damage.

## II. Introduction

Bone fragility is a major and growing public health burden, with approximately 37 million fragility fractures occurring annually worldwide in individuals aged over 55 [18], and the annual incidence of hip fractures alone projected to nearly double between 2018 and 2050 [19]. Much of this fragility traces to changes in geometry of cortical bone rather than changes in its material composition, as intracortical remodeling becomes unbalanced with age, vascular canals widen and coalesce, and whole-bone strength falls as roughly the seventh power of cortical porosity [20]. The hierarchical organization of bone and the variability of biological tissue make it difficult to study this geometric effect in isolation, so a controllable surrogate is useful. Calcium sulfate hemihydrate, established as both a bone-graft material and a brittle analog for cortical tissue, provides one. Plaster beams were drilled with defects, then repaired with the same plaster, and tested in three-point bending to determine how bulk flexural response changes as the number of repaired defects increases, and if the geometry of the resulting fracture path carries any mechanical information that defect count alone cannot.

### Cortical Bone Microstructure

The outermost structural layer of most bones is comprised of a dense osseous tissue called cortical bone[1]. It serves a protective role, providing rigid structural support for the rest of the bone[1]. Cortical bone is formed in a lamellar fashion with layers that are ∼5 μm thick[1, 2]. Within each layer, collagen fibers are parallel to each other, but collagen fiber orientation alternates between layers, contributing to bone’s strength[1, 2]. The inner and outer surfaces of cortical bone are comprised of circumferential lamellae, which separate cortical bone from other structures [1, 3]. In between circumferential lamellae, cortical bone is largely composed of osteons, with the spaces between them being filled by interstitial lamellae[1-3]. Osteons are the main structural unit of cortical bones and are up to 10 mm long and ∼200-400 μm wide[1, 2]. Osteons are enclosed by the cement line, which helps prevent crack propagation and separates osteons from interstitial and circumferential lamellae[1-3]. Osteons house many components that are critical to bone health, including blood vessels, lymphatic vessels, nerves, and osteocytes, which are connected via lacunae and a series of canals[1, 3].

### Bone Remodeling

Bone remodeling is a tightly regulated process that occurs across a lifetime, where old or damaged bone is replaced with new bone. This process is critical to maintain homeostasis and preserve the structural integrity of bone; it is performed by a basic multicellular unit (BMU), consisting of osteoclasts, osteoblasts, and the associated blood vessels[2, 4, 5]. Cortical bone specifically is remodeled through intracortical remodeling, which takes 4-6 months in healthy adults and is done in the following coordinated phases: activation, resorption, reversal, formation, and quiescence[5]. Intracortical remodeling is triggered and activation happens when osteocytes sense mechanical loading and microdamage to cortical bone[1, 5]. Osteocytes then regulate osteoclast and osteoblast activity relative to the nature and severity of the bone damage[1, 5]. During this resorption process, osteoclasts form a tunnel (called a cutting cone) that advances longitudinally down the bone’s axis, dissolving minerals and liberating collagen fragments[4, 5]. Once the collagen fragments are removed, the reversal phase happens, signaling the cease of resorption and the beginning of bone formation[5]. During formation, osteoblasts deposit concentric lamellae of new bone around a vascular channel, forming a mature osteon[2, 3, 5]. Cortical bone remodeling is vital for repairing bone damage and preventing damage accumulation[6, 7]. As humans age or contract bone diseases, intracortical remodeling can contribute to wider vascular channels and increased bone porosity, making the bone weaker [6, 7]. This highlights the need for continued research on how bone remodeling affects mechanical properties.

### Plaster as a Surrogate Material

Calcium sulfate hemihydrate (2CaSO_4_·H_2_O) commonly called Plaster of Paris is an inorganic salt produced from calcined gypsum and one of many forms of calcium sulfate. Historical evidence shows human use of plaster as far as 9000 years ago for creating structures and artwork, with the pieces found in the best condition being made of calcium sulfate hemihydrate produced using the same method we use today[8]. The use of calcium sulfate dihydrate to immobilize fractures was recorded as far back as the tenth century and became widely accepted by the nineteenth century[8]. In 1892 calcium sulfate dihydrate was successfully used as a bone grafting material to fill bone cavities associated with tuberculosis and has since been identified as one of the oldest biomaterials used for bone grafting[9-11]. Calcium sulfate dihydrate performs well clinically due to its fast degradation, predictable resorption rate *in vivo*, and complete resorption with minimal inflammatory response[10, 12, 13].

Calcium sulfate hemihydrate is produced in two forms, a (alpha) and ß (beta), with similar chemical properties and differing in size, lattice structure, and surface area[12]. In a process known as wet calcination, alpha-calcium sulfate hemihydrate is produced by calcining gypsum in an autoclave at 120^°^ C to 130^°^ C under steam pressure for five to seven hours[12]. Beta-calcium sulfate hemihydrate is produced through a process known as dry calcination by first milled and then heated to a temperature of 110^°^ to 130^°^ C[12]. Calcium sulfate hemihydrate shares characteristics close to natural bone, making it a suitable surrogate material for this study, with the drawback being that it is susceptible to brittle fractures and has lower mechanical stability[14].

### Three-Point Bending

Three-point bending is a common mechanical test used to assess the mechanical characteristics of a given material such as its flexural modulus, flexural strength, and flexural strain. In general three-point bending is performed by supporting a beam-like specimen at two ends while a force is applied at a midpoint with this specific study adapting three-point bending procedures from ASTM C1341-13[15]. Previous studies have used three point bending to investigate the fracture mechanisms of mouse bone, a commonly used model of human bone[16]. Brittle and quasi-brittle materials often fail from tensile cracking, making flexural tests such as three-point bending particularly useful for studying fracture behavior, flexural strength, and stiffness in materials such as plaster, ceramics, and bone analogs[17].

## III. Materials and Methods

### Sample Preparation

Specimens were fabricated from a standard formulation of calcium sulfate hemihydrate prepared at a 2:1 mass ratio of plaster to cold water. The components were combined and mixed until a homogenous mixture was obtained. The mixture was cast into silicone molds. Cast plates were allowed to cure under ambient laboratory conditions for 48 hours before demolding, and then sectioned into rectangular beams using a 12-inch hacksaw blade. Beams were progressively sanded by hand with abrasive papers of increasing grit, from 60 to 1200, to final dimensions of 5 mm x 15 mm x 90 mm, in accordance with ASTM C1341-13. A total of 38 specimens were produced, where 33 were reserved for experimental testing and 5 were used to calibrate the three point bending apparatus.

### Defect Introduction

Cylindrical through-thickness defects were introduced into each beam to produce controlled internal heterogeneities that were then repaired and tested as part of the whole specimen. On the upper face of each specimen, boundary lines were etched 15 mm from each end to define a 15 mm x 60 mm working region within which defects were placed. Defects were drilled using a Sherline Model 5410 tabletop CNC mill equipped with stepper-motor control on the X, Y, and Z axes and a 7/64-inch-double-sided drill bit. Each specimen was supported on a wooden backing block fixed at the center of the mill stage and clamped against the backing using a step-clamp and serrated wedge assembly secured by a T-slot nut. Defect coordinates were generated by a custom random-placement script constrained to the defined working region, and coordinates intersecting the workspace boundary and each other were discarded. Random placement was used to evaluate the bulk effect of the repaired-defects on a whole specimen rather than local effects of single defects at controlled locations. The coordinate sets were imported into FlashCut CNC for automated tool positioning, with the drill bit manually zeroed at the lower-left corner of the working region prior to each run. Each defect was drilled completely through the specimen thickness and into the underlying wooden block to ensure uniform geometry. Specimens were prepared in groups containing 0 through 10 defects, with triplicate specimens per group. All defects were recorded for subsequent analysis.

### Defect Repair

The underside of each specimen was sealed with painter’s tape to provide a backing surface for material infill. A batch of plaster was prepared at the same 2:1 mass ratio as before, with one drop of blue tissue marking dye (Medline SL662BL6-2/2236) added per batch to provide visual contrast between the original and repair material. The dyed mixture was loaded into a 3 mL syringe with the needle removed and was injected into each defect from the top surface until a positive meniscus formed at the opening. Filled specimens were cured under ambient conditions, after which underside tape was removed and excess repair material was sanded flush with 1000-grit abrasive paper until a sharp interface between the host plaster and the dyed infill was visible at both faces.

### Mechanical Testing

Three-point bending was performed on an Instron 5969 universal testing system following the procedure outlined in ASTM C1341-13. The support span was fixed at 80 mm, and each specimen was positioned across the supports with its mid-span aligned beneath the crosshead. The crosshead was advanced at a displacement rate of 0.21 mm/s and the test was continued until critical failure of the specimen. Time, crosshead extension, and load were sampled continuously throughout each test.

Raw load-extension records were processed through a custom Python pipeline. The point of first specimen-crosshead contact was identified as the earliest time at which the measured load rose above five standard deviations of the pre-contact baseline noise and remained above that threshold for at least five consecutive samples. The time, extension, and load axes were then rezeroed at that point. Flexural stress, ***σ***, and flexural strain, ***ε***, were computed from the rezeroed records using the standard three-point bending relations = 3FL2bd2 and = 6dL2, where F is the applied load, ***δ*** is the crosshead displacement, L = 80 mm is the support span, and b = 5 mm and d = 15 mm are the specimen width and depth, respectively. The maximum flexural stress and the corresponding flexural strain at peak stress were extracted directly from the stress-strain record. The flexural modulus, E, was obtained by linear regression of stress against strain over the 30 - 70 % range of the rising limb of maximum flexural stress, with the strain axis subsequently shifted by the intercept of the modulus fit so that the linear elastic region extrapolated through the origin. Work to failure was computed as the trapezoidal integral of stress with respect to strain from rezeroed contact to the peak.

### Fracture Tortuosity Analysis

The fracture surface of each specimen was photographed under consistent lighting to enable image-based quantification of fracture tortuosity. The fracture path in each image was extracted and analyzed using a custom Python pipeline built on the OpenCV, scikit-image, and SciPy libraries. Each image was first converted to grayscale and smoothed with a 5 × 5 Gaussian kernel to suppress surface texture, then segmented with an adaptive gaussian threshold (block size = 35, constant offset = 5) to isolate the dark fracture region from the surrounding intact material.

A morphological closing operation with a 3-pixel elliptical kernel was applied to bridge minor discontinuities in the segmented fracture, and connected components smaller than 50 pixels were removed to suppress residual noise. The largest remaining connected component was retained as the fracture trace.

The retained fracture region was reduced to a single-pixel-wide centerline by morphological skeletonization, and skeleton endpoints were identified through convolution with a 3 × 3 connectivity kernel. The geodesic path length, Lp, between the two terminal points was computed along the skeleton using Dijkstra’s algorithm with 8-connected neighborhoods and edge weights of 1 for orthogonal moves and 2 for diagonal moves. The straight-line Euclidean distance, Le, between the same two terminal points was computed directly from their image coordinates. Fracture tortuosity, **τ**, was then defined as the dimensionless ratio = LpLe with = 1 corresponding to a perfectly straight fracture and > 1 indicating a more deviated fracture path. Since tortuosity is a ratio of two lengths measured in the same image, no physical calibration of pixel size was required. For each specimen, the segmented fracture, the recovered skeleton, the endpoints, and the geodesic path were overlaid on the original image and archived for visual verification, and the computed values of geodesic path length, Euclidean distance, and tortuosity were exported to a results table.

## IV. Results

### Flexural Response and Stress-Strain Behavior

All 33 experimental specimens failed in a brittle manner, with the load rising approximately linearly to a single peak followed by an abrupt drop, consistent with previously reported behavior of calcium sulfate dihydrate under flexure. The linearized modulus fits achieved coefficients of determination R2 0.984 across all 33 specimens, indicating that the 30 - 70 % peak-stress fit window captured the elastic regime cleanly in each test.

**Figure 1.**
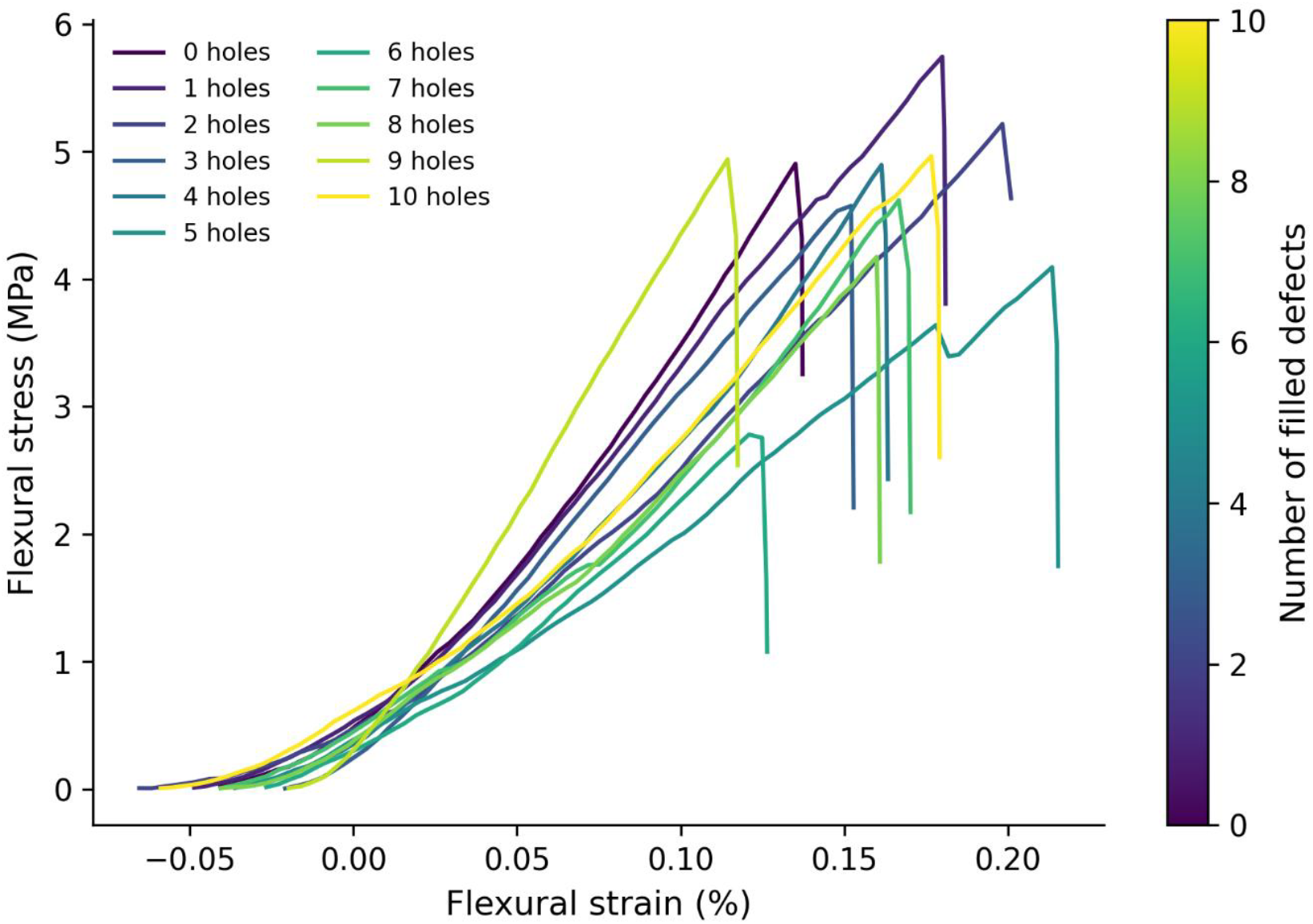
Representative flexural stress–strain curves for plaster beams containing 0 to 10 repaired defects. One representative specimen per defect group is plotted. All specimens failed in a brittle manner, with the load rising approximately linearly to a single peak followed by an abrupt drop. Stress was computed from the re-zeroed load–extension record using the standard three-point bending relation σ = 3FL/(2bd^2^), with L = 80 mm, b = 5 mm, and d = 15 mm; strain was computed as ε = 6δd/L^2^.

### Effect of Defect Count on Mechanical Properties

Across the range of 0 to 10 filled defects, the mean flexural strength varied between 3.25 1.07 MPa and 5.93 0.39 MPa, with the unmodified control specimens producing 4.86 0.36 MPa. A linear regression of maximum flexural stress against defect count yielded a slope of -0.089 MPa per defect (R2 = 0.08, p = 0.10), and the corresponding Spearman rank correlation was ρ = -0.28 (p = 0.12), indicating only a weak and statistically non-significant monotonic decline in strength as the number of repaired defects increased.

The flexural modulus, E, was unaffected by defect count. Group means ranged from 2.32 0.38 GPa to 3.88 0.86 GPa with no trend (slope = 16.7 MPa per defect, R2 = 0.01, p = 0.64; Spearman ρ = 0.004, p = 0.98). Strain at peak stress showed no significant dependence on defect count (slope = -0.0035% per defect, R2 = 0.12, p = 0.050).

Work to failure decreased significantly with increasing defect count. Group means fell from 4.17 0.70 × 10-3 MJ/m3 in the control specimens to 2.59 1.85 × 10-3 MJ/m3 at 6 defects, with a linear slope of -1.49 × 10-4 MJ/m3 per defect (R2 = 0.15, p = 0.028; Spearman ρ = -0.38, p = 0.028). The quadratic fit to the same data produced a marginal improvement in variance (R2 = 0.20), suggesting a predominantly linear monotonic degradation in energy absorption as defect density increased.

**Figure 2.**
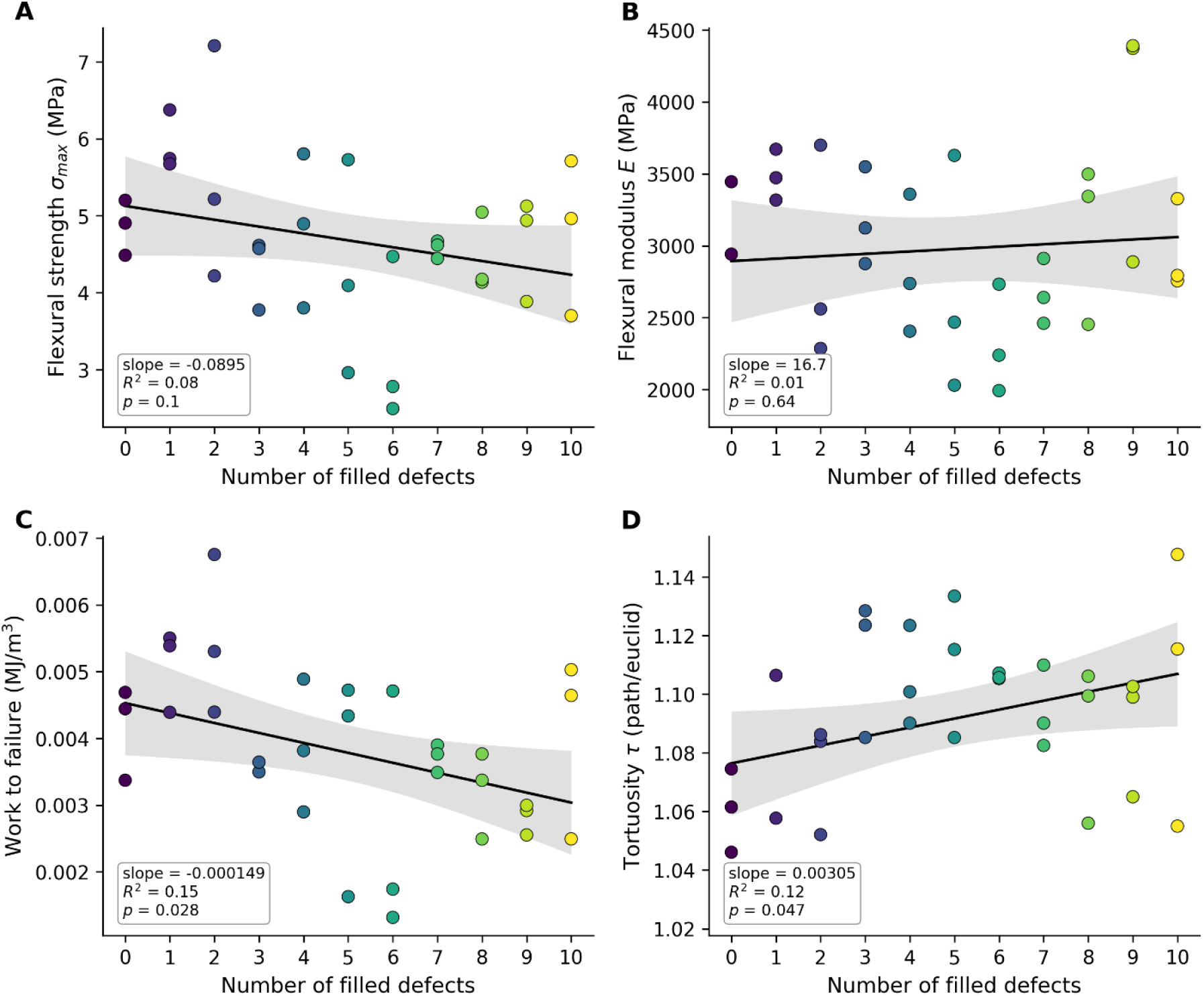
Per-specimen mechanical and geometric properties plotted against the number of repaired defects (n = 3 per group, n = 33 total). (A) Flexural strength σmax. (B) Flexural modulus E. (C) Work to failure Wf. (D) Fracture-path tortuosity τ. Each point represents one specimen, colored by defect count. Solid lines show ordinary-least-squares linear regressions and shaded bands show 95 % confidence intervals about the fit.

### Qualitative Fracture-Path Observations

Due to the random generation of defect coordinates, only a subset of induced defects fell within the high-tensile-stress zone immediately beneath the crosshead, and the remaining defects were located outside of failure-relevant regions. Across specimens in which one or more defects did intersect the failure path, the propagating fracture was consistently observed to deflect around the perimeter of the repair fill rather than transect it. In these instances, the fracture surface curved along the host-repair boundary and re-emerged into the host matrix on the far side of the defect, leaving the infill intact.

### Fracture Tortuosity

Fracture path tortuosity, **τ**, quantified by skeletonization of post-test fracture images, increased modestly but significantly with defect count. The group mean rose from = 1.061 0.014 in the control specimens to = 1.106 0.047 at 10 defects, and the linear regression yielded a slope of 0.0031 per defect (R2 = 0.12, p = 0.047; Spearman ρ = 0.31, p = 0.082). This corresponds to a roughly 4 % increase in cumulative fracture-path length over the tested defect range, indicating that crack trajectories became progressively more deviated as more repair interfaces were present to be encountered.

Linear regression showed a significant negative relationship between tortuosity and flexural strength (slope = -13.8 MPa per unit, R2 = 0.15, p = 0.025; Spearman ρ = -0.43, p = 0.013), with weaker non-significant trends for modulus (R2 = 0.09, p = 0.089) and work to failure (R2 = 0.06, p = 0.17). The tortuosity-max flexural stress correlation was of a greater magnitude than the corresponding defect count-max flexural stress correlation, with both a larger Spearman coefficient (0.43 vs 0.28) and a lower p-value (0.013 vs 0.12).

**Figure 3.**
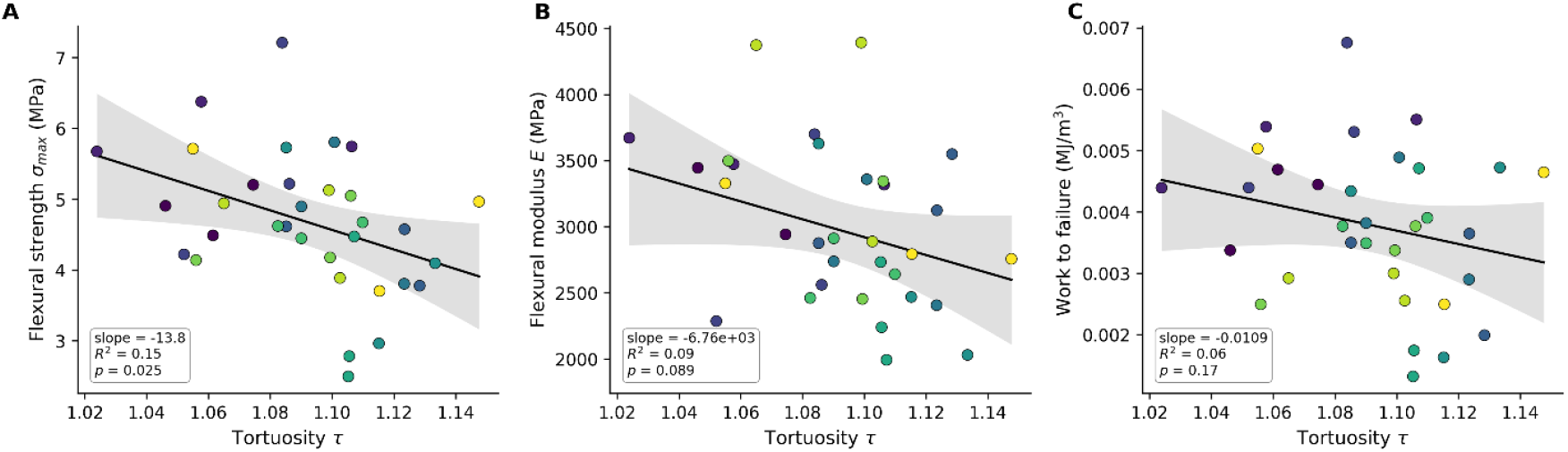
Mechanical properties plotted against fracture-path tortuosity τ across all 33 experimental specimens. (A) Flexural strength σmax. (B) Flexural modulus E. (C) Work to failure Wf. Each point represents one specimen, colored by defect count using the same scale as Figure 2. Solid lines show ordinary-least-squares linear regressions and shaded bands show 95 % confidence intervals about the fit.

## V. Discussion

The principal findings are that flexural strength and modulus were only weakly and non-significantly reduced by the addition of up to ten repaired defects, that the energy absorbed prior to failure declined significantly with defect count, that cracks encountering a repair within the failure zone consistently deflected around the perimeter of the repair interface rather than transecting it, and that fracture tortuosity rose significantly with defect count and was itself a stronger predictor of strength than defect count alone.

The qualitative observation that fractures deflected around the repaired defects rather than through them is the central mechanistic result and frames the rest of the quantitative findings. The repaired defect itself must be at least as fracture-resistant as the surrounding host plaster, since if it were weaker, the crack would have preferentially transected the repair on its straight-line path rather than detouring around it. The most likely explanation is that the repair plaster, injected wet into a dry confining cavity, cured under partial restraint and against a roughened interior wall, producing a locally denser fill than the host material, which was cast against smooth silicone with one free upper surface. The fact that the deflected crack tracked the host– repair boundary itself, rather than passing through the host matrix on a path that happened to miss the repair, indicates that the interface is a preferential weak path. The repair sites therefore behave as a two-component system, consisting of a tough repair bonded to the host through a comparatively weaker interfacial region. This combination of stiff inclusions surrounded by weak interfaces is qualitatively analogous to the cement-line architecture surrounding osteons in cortical bone, where cracks have been shown to deflect preferentially along the cement line while leaving the osteonal interior largely intact.

The significant negative correlation between tortuosity and flexural strength (R^2^ = 0.15, p = 0.025; ρ = −0.43, p = 0.013), and the fact that this correlation is stronger than the relationship between defect count and strength (ρ = −0.28, p = 0.12), is the most informative quantitative finding of the study. We interpret this difference in predictive power because of the random defect placement protocol. Defect count, as a specimen-level descriptor, treats every introduced hole as mechanically equivalent regardless of where it was located within the beam. However, only the subset of defects that intersected the failure path could have influenced the fracture outcome, and the remainder were mechanically inert with respect to the test. Tortuosity, by contrast, is computed from the realized crack trajectory and therefore registers a contribution only from defects that engaged with the propagating crack. In this sense, τ functions as a measure of effective defect interaction rather than nominal defect density, which explains why it correlates with σmax more strongly than the raw defect count does. This interpretation suggests that, in studies of heterogeneous brittle materials where defect placement cannot be precisely controlled, post-mortem geometric descriptors of the fracture path may offer a more direct estimator of the mechanical impact of the defect population than counts or densities computed from the as-built geometry.

## VI. Conclusion

Flexural response and fracture path tortuosity of plaster-of-Paris beams containing 0 to 10 cylindrical defects that were drilled and subsequently repaired with the same plaster formulation were quantified. Across the 33 specimens tested in three-point bending with ASTM C1341-13, the bulk flexural strength and modulus showed only a weak and statistically non-significant dependence on the number of repaired defects, suggesting that the restoration achieved by the repair process preserved the principal load-bearing characteristics of the beam. The work to failure and the in-plane fracture tortuosity, however, both varied significantly with defect count. Qualitative inspection of fracture surfaces revealed that the propagating fracture consistently deflected around the perimeter of the repaired defect along the host-repair interface rather than transecting the repair material itself, which is a deflection geometry analogous to the cement-line-mediated fracture redirection observed in cortical bone, but here isolated from the additional plastic-dissipation mechanisms that occur in living tissue. A direct regression of strength against tortuosity revealed a stronger negative correlation than was obtained against defect count alone. This difference could reflect that tortuosity measures effective defect interaction with the fracture path, and defect count treats all defects as mechanically equivalent regardless of their positions relative to the failure zone. This suggests that calcium sulfate dihydrate could be used as a platform for isolating the geometric contributions of repaired defects to fracture behavior, and identifies fracture-path tortuosity as a low-cost, image-based metric that captures aspects of mechanical impact of internal heterogeneity that defect counts do not.

